# Cohesin residence time gates 3D genome response to histone hyperacetylation

**DOI:** 10.64898/2026.07.01.735920

**Authors:** Rebecca G Smith, Kathleen L Schiela, Hannah M Wilson, Ryan A Williams, James Johnson, Carson B Cohen, Wei-Ting Yueh, Amy M Whitaker, Neil Johnson, Masato T Kanemaki, Yu Liu

## Abstract

Cohesin-mediated loop extrusion and chromatin state-dependent compartmentalization are major drivers of three-dimensional (3D) genome organization. Although epigenomic perturbations are widely assumed to reshape chromatin architecture, the mechanisms that determine how changes in chromatin state are translated into structural reorganization remain poorly understood. Here, we identify cohesin residence time as a key regulator of the genome’s architectural response to histone hyperacetylation induced by histone deacetylase inhibition (HDACi). Acute depletion of RAD21 or CTCF weakens chromatin loops but preserves HDACi-induced changes in compartmentalization, contact-scaling behavior, and loop density. In contrast, perturbation of cohesin loading or release produces opposing effects: NIPBL depletion sensitizes and amplifies architectural responses to HDACi, whereas WAPL loss renders the genome largely refractory to HDACi-induced remodeling, suppressing changes in compartments and loop density while stabilizing CTCF-anchored loops. These distinct architectural outcomes occur despite comparable levels of HDACi-induced histone hyperacetylation across genotypes, indicating that differential epigenomic input is not responsible for the observed effects. Together, our findings demonstrate that dynamic cohesin turnover, rather than cohesin chromatin association alone, governs whether epigenomic perturbations are converted into higher-order genome reorganization. These results establish cohesin residence time as a molecular gate linking chromatin state to 3D genome architecture and reveal a previously unrecognized principle underlying chromatin architecture plasticity.

## Introduction

The cohesin complex organizes the interphase genome through loop extrusion, a process driven by a dynamic cycle of cohesin loading by NIPBL and release by WAPL (Fig. 1A) (Sanborn et al. 2015; Fudenberg et al. 2016; Haarhuis et al. 2017; Rao et al. 2017; Schwarzer et al. 2017). Perturbations to this cycle alter cohesin residence time on chromatin and reshape genome architecture: loss of NIPBL weakens loops and enhances compartmentalization, whereas loss of WAPL stabilizes loops and attenuates compartment strength (Haarhuis et al. 2017; Schwarzer et al. 2017; Wutz et al. 2017). Loop extrusion is halted at CTCF-binding sites, where stalled cohesin establishes topologically associating domain (TAD) boundaries and stable chromatin loops; accordingly, CTCF depletion abolishes these structures (Sanborn et al. 2015; Fudenberg et al. 2016; Nora et al. 2017). Together, these observations establish cohesin dynamics as a central determinant of 3D genome organization.

**Fig. 1.**
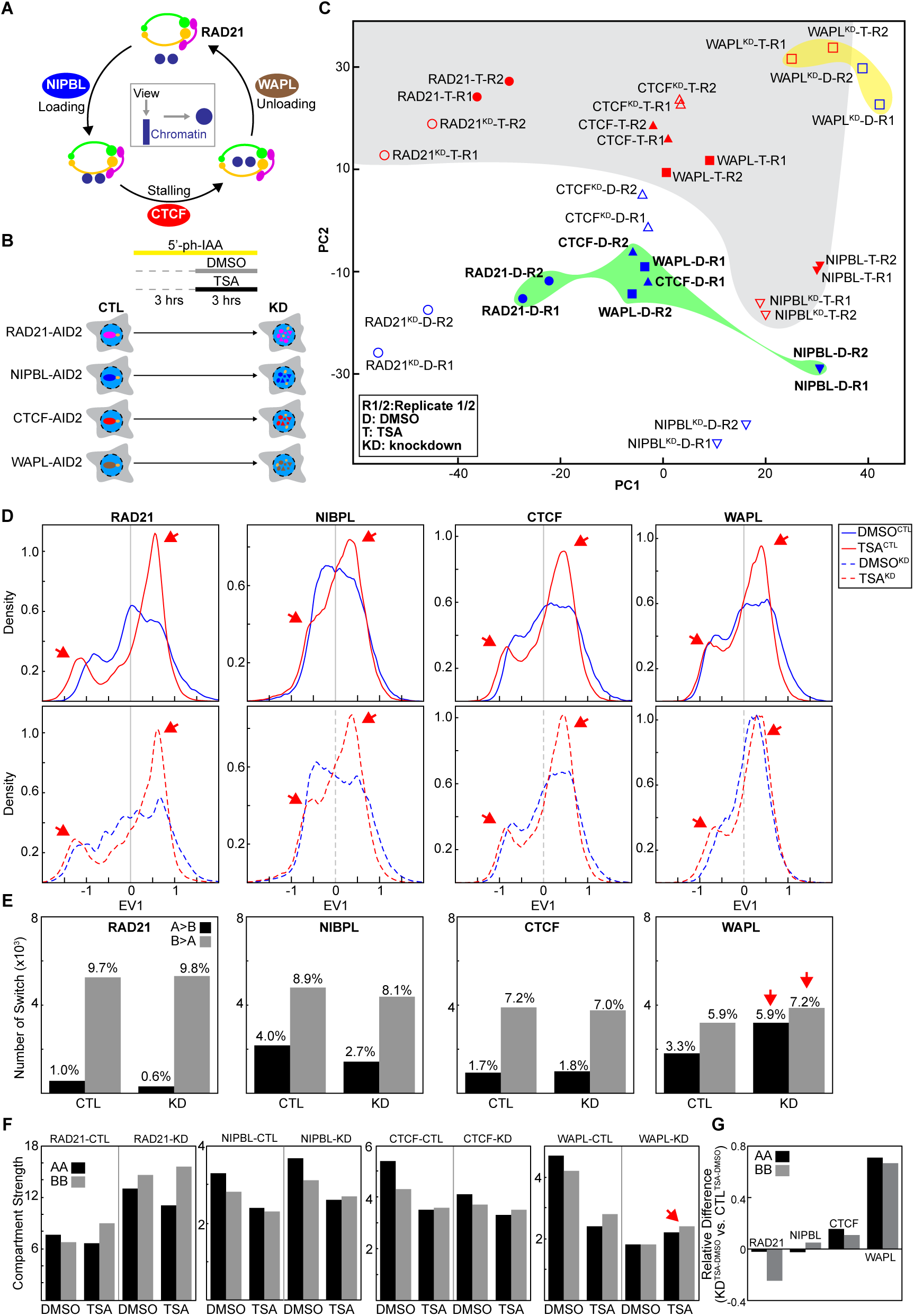
Influence of cohesin dynamics on genome architectural changes following TSA treatment. **(A)** Schematic of the cohesin regulatory cycle. NIPBL loads cohesin onto chromatin, CTCF stalls extruding cohesin to define loop and boundary positions, and WAPL promotes cohesin release. **(B)** Experimental design illustrating the systematic depletion of cohesin subunits or regulators to assess their roles in genomic structure responses to TSA-induced histone hyperacetylation. **(C)** Prinicipal component analysis (PCA) of eigenvector 1 values (EV1), demonstrating compartment profile of EV1 in each condition. CTL and KD indicates non-treatment or treatment of 5’-Ph-IAA, which incudes degradation of targeted gene. D and T represent DMSO- and TSA-treatment, and R1/R2 represents biological replicates 1 and 2, respectively. The green shading highlights the variation between CTL, DMSO-treated cells, while the grey shading shows the grouping of all TSA-treated cell lines, and the yellow shading shows the WAPL-KD cells with DMSO or TSA treatments. **(D)** Compartment (A/B) patterns in DMSO- and TSA-treated cells across RAD21, NIPBL, CTCF, and WAPL degron backgrounds. For each degron line, the non-5’-Ph-IAA condition serves as the matched CTL, and the 5’-Ph-IAA-treated condition represents the acute depletion (KD), as denoted by solid and dashed lines. Blue and red represent DMSO- and TSA-treatment. **(E)** Compartment switching ratios (A-to-B and B-to-A) comparing DMSO- versus TSA-treated CTL and KD cells across the same backgrounds shown in **(D). (F)** Compartment strength of A or B compartments comparing DMSO and TSA treatment for CTL (non-IAA) and KD (IAA) conditions across the same backgrounds as in **(D)**. **(G)** Relative difference of AA and BB compartment strength changes upon TSA treatment between CTL and KD conditions (see methods).

Chromatin state is also closely linked to genome folding. Histone deacetylase inhibitors (HDACis) induce widespread histone hyperacetylation and have been shown to weaken nucleosome interactions, remodel chromatin architecture, and alter nuclear organization (Nozaki et al. 2017; Sanders et al. 2022; Paldi F 2024; Smith et al. 2025). However, the mechanisms that determine how epigenomic perturbations are translated into changes in genome architecture remain poorly understood. In our recent work, we found that histone hyperacetylation selectively weakens cohesin-mediated interactions within TADs while largely preserving global cohesin occupancy on chromatin and maintaining CTCF-anchored loops(Smith et al. 2025). These findings suggested that the architectural response to histone hyperacetylation is governed not simply by cohesin occupancy, but by the manner and duration with which cohesin engages chromatin. Since cohesin residence time is a principal determinant of cohesin-chromatin interactions, we hypothesized that it may act as a key regulator of genome architectural responsiveness to epigenomic change.

To test this hypothesis, we combined acute perturbations of the cohesin regulatory cycle with HDAC inhibition to determine how cohesin dynamics influence structural responses to histone hyperacetylation. We find that depletion of cohesin itself (RAD21) or removal of boundary elements (CTCF) leaves HDACi-induced changes in chromatin compartmentalization, contact-scaling behavior, and loop density largely intact. In contrast, manipulating cohesin residence time produces opposing outcomes: loss of NIPBL sensitizes genome architecture to HDACi-induced remodeling, whereas loss of WAPL suppresses these responses and further stabilizes CTCF-anchored loops. Together, our findings identify cohesin residence time as a molecular gate that determines whether epigenomic perturbations are translated into large-scale reorganization of the 3D genome, revealing a previously unrecognized principle linking chromatin state to genome architecture.

## Results and Discussion

To investigate how cohesin dynamics influence genome architectural responses to histone hyperacetylation, we utilized second-generation HCT-116 auxin-inducible degron (AID2) cell lines, which enable rapid depletion of target proteins(Yesbolatova et al. 2020). Cells were treated with 5′-Ph-IAA for three hours to induce degradation of RAD21, CTCF, NIPBL, or WAPL, followed by treatment with the histone deacetylase inhibitor trichostatin A (TSA) (Figure 1B). Cells were subsequently fixed and analyzed by Hi-C(Lieberman-Aiden et al. 2009). Because acute RAD21 depletion induces G2/M arrest, G1-phase cells were isolated by flow cytometry prior to Hi-C analysis(Liu and Dekker 2022). Throughout this study, cells treated with 5′-Ph-IAA are referred to as knockdown (KD) samples, whereas untreated cells serve as matched controls (CTLs).

Western blot analysis confirmed efficient degradation of RAD21, CTCF, NIPBL, and WAPL following 5′-Ph-IAA treatment. TSA induced comparable levels of histone hyperacetylation in all genetic backgrounds, indicating that cohesin pathway perturbations do not alter the primary epigenomic response to HDAC inhibition (Supplemental Fig. S1A, two biological replicates). Furthermore, cell-cycle profiles remained largely unchanged following target depletion, TSA treatment, or their combination (Supplementary Fig. S1B). These results establish that the differential architectural responses described below cannot be explained by differences in histone hyperacetylation or cell-cycle composition.

Since chromatin compartmentalization is closely associated with transcriptional activity and cell identity(Dixon et al. 2012; Rao et al. 2014), we first assessed global compartment differences across all samples using principal component analysis (PCA) of the first eigenvector (EV1) (See Methods). Although all degron cell lines were derived from the parental HCT-116 line, they exhibited distinct compartment profiles(Fig. 1C, green area, two biological replicates per condition). CTCF- and WAPL-degron cells clustered closely together, whereas RAD21-degron cells formed a separate group. Notably, NIPBL-degron cells were clearly separated from all other cell lines. This observation may reflect the established role of NIPBL in transcriptional regulation (Dorsett and Krantz 2009; Zuin et al. 2014), although clonal variation introduced during cell line generation cannot be excluded. Importantly, these baseline differences do not affect our analysis, which focuses on TSA-induced architectural changes within each matched CTL-KD pair.

To evaluate the impact of HDAC inhibition on chromatin compartmentalization, we compared compartment profiles before and after TSA treatment. Consistent with our previous observations, TSA induced substantial compartment remodeling in CTL cells. However, the magnitude of these changes varied among cohesin perturbation backgrounds. Most notably, WAPL depletion produced the weakest TSA-induced compartmental response, suggesting that prolonged cohesin residence time buffers genome architecture against histone hyperacetylation (Fig. 1C).

We next examined compartment organization by analyzing EV1 distributions, after orienting EV1 such that positive and negative values correspond to A and B compartment identities, respectively(Lieberman-Aiden et al. 2009). In CTL cells, TSA treatment induced a pronounced bimodal EV1 distribution, indicating increased polarization of chromatin regions toward either A or B compartment states (Fig. 1D and Supplemental Fig. S1C). This TSA-induced bimodality was observed across all CTL backgrounds, although it was less pronounced in NIPBL-degron cells, consistent with its distinct baseline compartment profile (Fig. 1C).

Acute depletion of RAD21, NIPBL, or CTCF altered baseline EV1 distributions but did not abolish the TSA-induced shift toward a bimodal compartment state. In contrast, WAPL depletion markedly blunted this response, with TSA inducing only limited additional EV1 redistribution (Fig. 1E and Supplemental Fig. S1D). Consistent with previous reports (Wutz et al. 2017), WAPL depletion alone increased the density of positive EV1 values while shifting both positive and negative EV1 distributions toward zero, indicative of reduced compartment segregation. Together, these findings suggest that prolonged cohesin residence time limits HDACi-induced redistribution of compartment identity.

We next quantified compartment switching following TSA treatment. Depletion of RAD21, NIPBL, or CTCF largely preserved TSA-induced A-to-B and B-to-A transitions. In contrast, WAPL depletion markedly increased compartment switching events, particularly A-to-B transitions (Fig. 1F and Supplemental Fig. S1E). This increase may reflect reduced compartment segregation in WAPL-depleted cells, thereby lowering the barrier to compartment transitions. Thus, prolonged cohesin residence time suppresses global redistribution of compartment identity in response to histone hyperacetylation while increasing local compartment switching events.

We then assessed compartment strength by quantifying AA and BB compartment interactions. In CTL cells, TSA treatment generally reduced AA and BB interactions, with the exception of RAD21 CTL cells, in which BB interactions increased following TSA treatment (Fig. 1G and Supplemental Fig. S1F). As our analysis compares TSA-treated and untreated samples within each matched genetic background, these baseline differences do not affect the interpretation of TSA-induced responses.

Acute depletion of RAD21, NIPBL, or CTCF largely preserved TSA-induced changes in compartment strength. In contrast, WAPL depletion markedly attenuated TSA-induced compartment remodeling. Quantitative comparison of TSA-induced changes in AA and BB interactions between CTL and KD cells confirmed that WAPL loss produced the strongest suppression of architectural responsiveness to TSA (Fig. 1H and Supplemental Fig. S1G). Collectively, these results demonstrate that prolonged cohesin residence time constrains both compartment identity redistribution and compartment-strength remodeling in response to histone hyperacetylation.

We next examined how cohesin dynamics influence local chromatin folding following histone hyperacetylation. Consistent with our previous study(Smith et al. 2025), TSA weakened local chromatin interactions within TADs, evident as reduced contact frequencies (blue) in differential Hi-C maps (Fig. 2A and Supplemental Fig. S2A). Acute depletion of RAD21 or NIPBL further enhanced this reduction, whereas depletion of CTCF or WAPL attenuated it (Fig. 2B and Supplemental Fig. S2B). Notably, perturbation of cohesin turnover through NIPBL or WAPL depletion produced substantially larger effects than depletion of RAD21 or CTCF, indicating that cohesin dynamics are a stronger determinant of HDACi responsiveness than cohesin abundance or boundary insulation. Scaling analysis further supported this conclusion. Consistent with our previous observations(Smith et al. 2025), TSA preferentially altered short-range chromatin interactions in CTL cells, producing characteristic changes in contact probability scaling curves. Acute depletion of RAD21, NIPBL, or CTCF did not abolish these TSA-induced scaling changes (Fig. 2C and Supplemental Fig. S2C). Direct comparison of TSA-induced scaling responses between CTL and KD cells demonstrated minimal effects of RAD21 or CTCF depletion. In contrast, WAPL depletion largely abolished TSA-induced scaling changes, whereas NIPBL depletion markedly amplified them (Fig. 2D and Supplemental Fig. S2D). These opposing phenotypes demonstrate that cohesin residence time, rather than cohesin abundance alone, determines the sensitivity of chromatin folding to histone hyperacetylation.

**Fig. 2.**
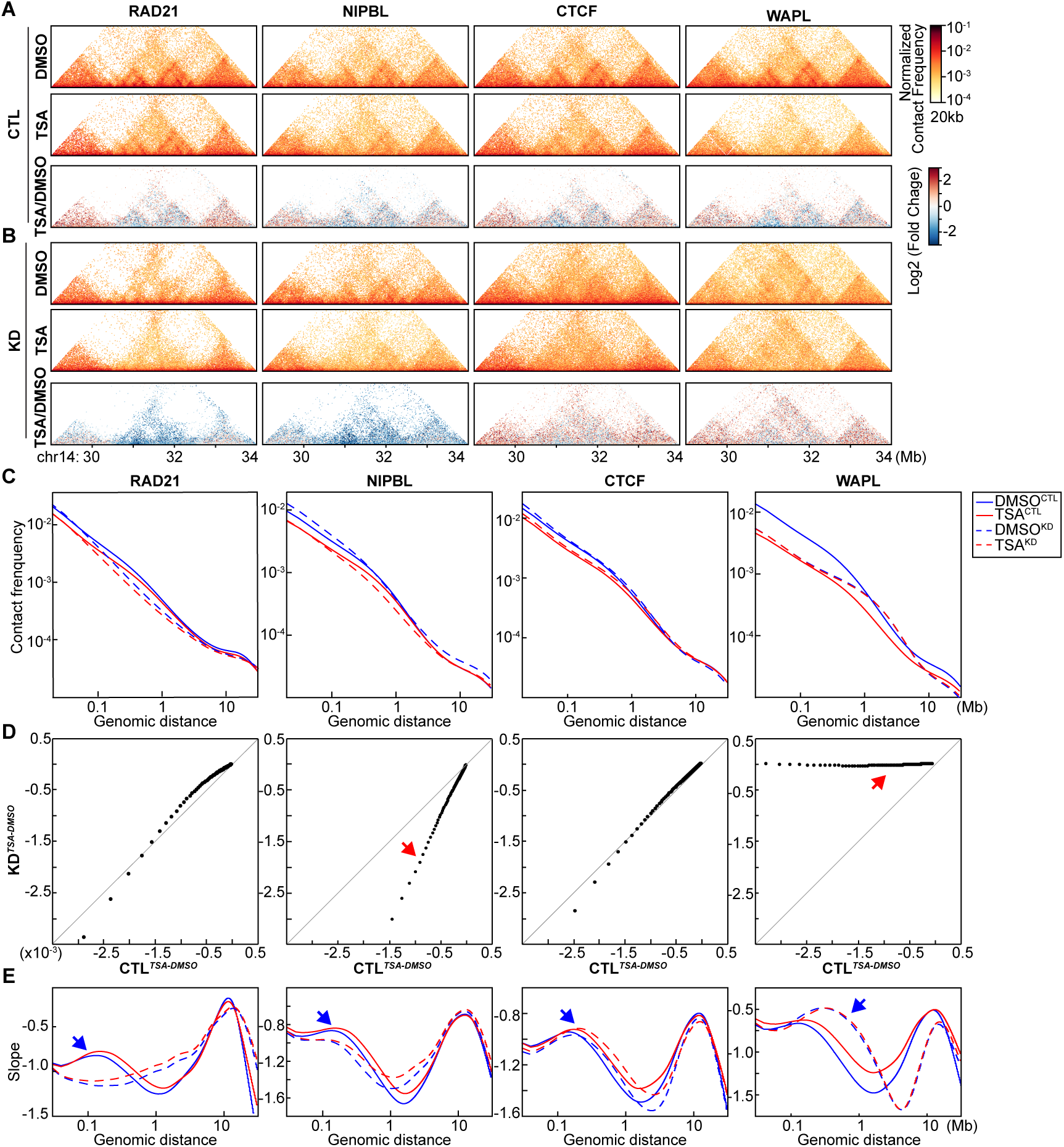
Altering cohesin dynamics produces opposing effects on TSA-induced scaling changes. **(A) and (B)** Hi-C interaction maps for each each CTL and KD cells treated with DMSO or TSA. Log2 fold change heat maps show the difference of chromatin interactions between TSA- and DMSO-treated cells within each condition. Data for the 29-24 Mb region on chromosome 14 are shown. **(C)** *P(s)* plots from Hi-C data of cells treated with DMSO or TSA, in the presence or absence of the indicated cohesin components. Blue and red lines represent DMSO- and TSA-treated cells, respectively; solid and dashed lines denote presence or absence of cohesin components, respectively. **(D)** Difference in scaling plots illustrating the effect of histone hyperacetylation in the presence (x) versus absence (y) of the indicated cohesin components. Red arrows highlight the difference between CTL and KD. **(E)** Derivatives of each corresponding P(s) plot from Hi-C data of cells treated with DMSO or TSA, without or without protein degradation. Blue arrows mark the peaks corresponding to cohesin-mediated loops observed in DMSO-treated cells.

We next quantified cohesin loop density using derivative analysis. Consistent with our previous findings(Smith et al. 2025), TSA reduced cohesin loop density in all CTL backgrounds. As expected, acute depletion of RAD21 eliminated cohesin-mediated loops(Rao et al. 2017; Wutz et al. 2017; Liu and Dekker 2022). Depletion of NIPBL or CTCF did not substantially alter the TSA-induced reduction in loop-associated signals. Strikingly, WAPL depletion almost completely prevented TSA-induced reductions in cohesin loop density (Fig. 2E and Supplemental Fig. S2E), indicating that prolonged cohesin residence time stabilizes cohesin-mediated loop interactions against histone hyperacetylation.

We next investigated whether cohesin dynamics similarly regulate stable CTCF-anchored loops. Aggregate Hi-C analysis of 3,165 CTCF-CTCF loops revealed that TSA modestly reduced loop strength in CTL HCT-116 cells, as measured by aggregate loop maps and loop-line fold changes, although this reduction was less pronounced in RAD21 CTL cells (Fig. 3A and Supplemental Fig. S3A). This differs from our previous observations in HAP1 cells, where CTCF-CTCF loops were largely insensitive to TSA treatment(Smith et al. 2025), suggesting either cell-type-specific differences or modest effects associated with the degron-engineered cell lines.

**Fig. 3.**
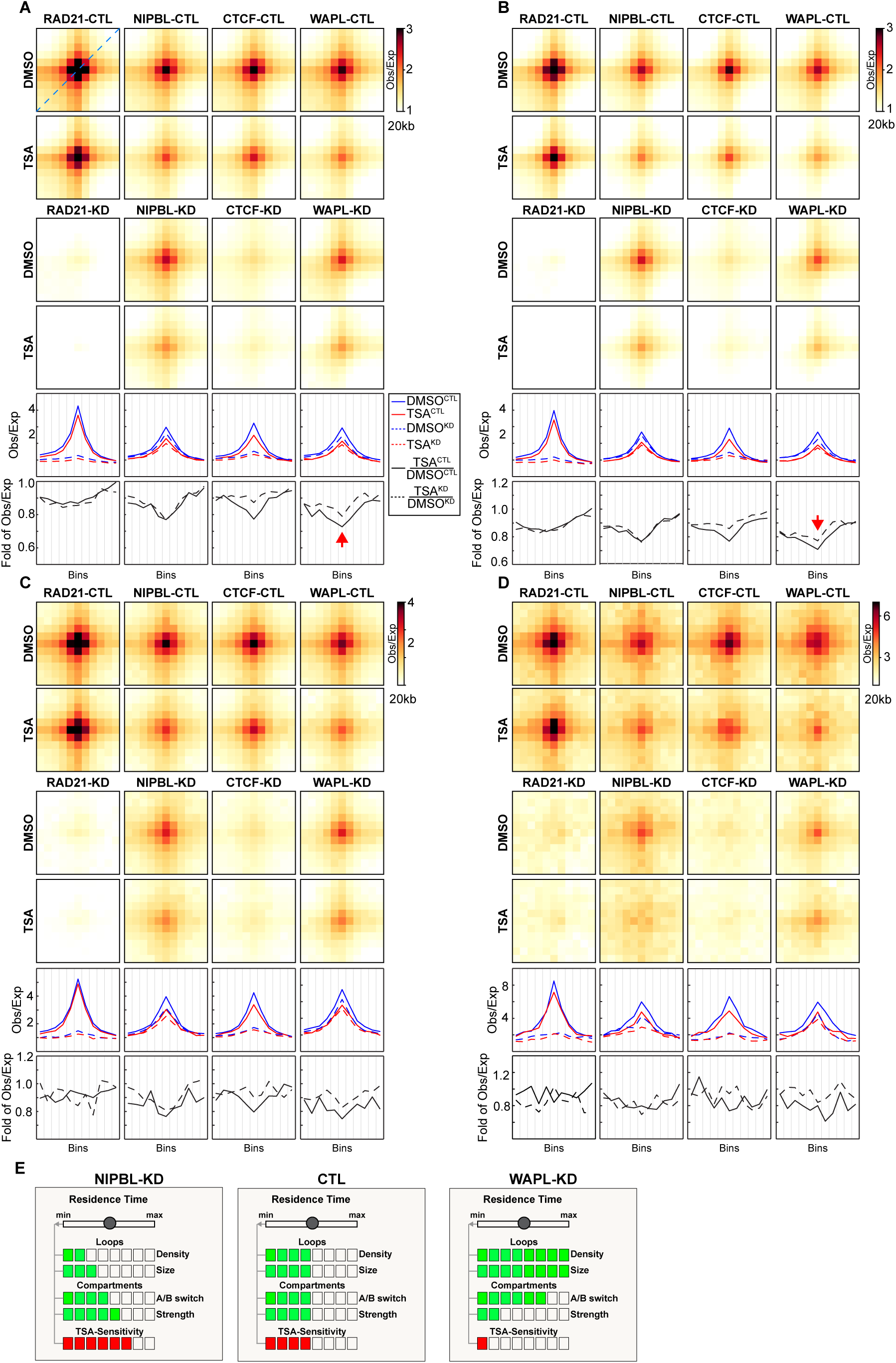
WAPL loss suppressed TSA-induced loop remodeling. **(A)** Aggregated Hi-C data at 3,165 CTCF-CTCF loops identified in HCT-116 cells according to(Rao et al. 2017). The top two rows show loop aggregate heatmaps for DMSO- or TSA-treated control cells; The middle rows show corresponding aggregates for DMSO- or TSA-treated cells following acute depletion of the indicated cohesin components and regulators. Loop strength was quantified along the diagonal from the bottom-left corner to the top-right of each aggregate heatmap (the “loop line”), illustrated by the blue diagonal in the top-left panel in **(A)**. The fifth row displays loop line profiles for each degron background, with blue and red lines representing DMSO- and TSA-treated samples, respectively. Solid and dashed lines denote non-5-Ph-IAA and 5-Ph-IAA treated conditions. The bottom row shows the TSA/DMSO fold-change in loop-line signal for CTL (solid lines) and degron (dashed lines) cells, indicating how depletion of specific cohesin regulators alters HDACi-induced loop remodeling. **(B)**-**(D)** Aggregated Hi-C data of different sizes of CTCF-CTCF loops. **(B)**, 100-500kb; **(C)**, 0.5-1mb; **(D)**, greater than 1mb. Red arrows highlight representative loop classes where TSA-induced remodeling in CTL cells is diminished upon CTCF or WAPL loss. **(E)** A schematic representing how cohesin residence time influences chromatin loop density and size, compartment patterns and strength, and subsequently, chromatin’s structural sensitivity to TSA treatment. Increased cohesin residence time on chromatin leads to resistance of structural changes induced by TSA, while decreased residence time renders chromatin more sensitive to structural perturbation.

Acute perturbation of cohesin regulators revealed distinct mechanisms underlying these responses. As expected, depletion of RAD21 or CTCF abolished CTCF-anchored loops regardless of TSA treatment. Consequently, TSA-induced changes were largely absent in these backgrounds because the loop structures themselves had already been eliminated. In contrast, WAPL depletion robustly prevented TSA-induced weakening of CTCF-CTCF loops, as demonstrated by both aggregate loop maps and loop-line fold changes (Fig. 3A and Supplemental Fig. S3A). This protective effect extended across all loop size classes (Fig. 3B-D and Supplemental Fig. S3B-D), indicating that prolonged cohesin residence time globally stabilizes CTCF-mediated chromatin architecture against epigenomic perturbation.

Importantly, these phenotypes could not be attributed to differential histone hyperacetylation, which was comparable across all degron backgrounds (Supplemental Fig. S1A). Thus, altered architectural responsiveness, rather than differences in the epigenomic input, accounts for the observed structural phenotypes.

Collectively, these findings establish cohesin turnover as the molecular gate that determines whether epigenomic perturbations are translated into structural remodeling of the genome. Although histone hyperacetylation provides a common epigenomic signal, its architectural consequences depend on the dynamic interaction of cohesin with chromatin. Perturbing cohesin abundance alone has relatively modest effects on HDACi-induced genome remodeling, whereas altering cohesin residence time fundamentally changes the architectural response. Prolonged cohesin residence time, enforced by WAPL depletion, stabilizes chromatin architecture and renders it refractory to epigenome-driven remodeling, whereas reduced cohesin residence time, caused by NIPBL depletion, sensitizes chromatin folding to histone hyperacetylation (Fig. 4). We therefore propose that cohesin residence time constitutes a fundamental architectural gate that determines whether epigenetic perturbations are translated into changes in 3D genome organization, providing a unifying framework for the context-dependent architectural responses to histone acetylation observed across different cellular states.

## Acknowledgements

We thank all the members of the Liu laboratory for discussion. We thank the assistance and support from the Biostatistics and Bioinformatics Facility, Genomic Resources Facility, Cell Sorting Facility, and Cell Culture Facility at Fox Chase Cancer Center. Research reported in this publication was supported by the National Cancer Institute of the National Institutes of Health under Award Number P30CA006927. We acknowledge support from the National Institute of General Medical Science (R35GM154879 to Y.L. and R35GM155098 to A.M.W.) and W. W. Smith Charitable Trust Grant (C2407 to Y.L. and C2302 to A.M.W.).

## Author Contribution Statement

Y.L. conceived and designed the project. R.G.S., K.L.S., and H.M.W. performed all the experiments with assistance of C.B.C and W.Y. and advice of Y.L., A.M.W. and N.J. M.T.K. generated all degron cells. R.G.S. processed and analyzed all the data with assistance of R.A.W, K.L.S and H.M.W. Y.L. supervised the project and Y.L. wrote the manuscript with assistance of R.G.S.

## Competing Interests Statement

The authors have no competing interests.

## Conflicts of interest

There is no conflict of interest.

## Materials and Methods

### Cell culture and chemicals

HCT-116-mAID-RAD21 degron2 cells were kindly provided by Yesbolatova, et al, 2020. These cells were cultured in McCoy’s 5A medium, GlutaMAX supplement (Gibco, 36600021) supplemented with 10% FBS (Gibco, 16000044) and 1% penicillin-streptomycin (Gibco, 15140) at 37°C in 5% CO2. Trichostatin A (#T8552) and 5’-ph-IAA (SML3574) were purchased from Sigma-Aldrich, USA and dissolved in DMSO (#D2650) before treating the cells. Degron cell lines were incubated with either DMSO or 5’-Ph-IAA at 37°C for 6 hours, with DMSO or TSA being added at the 3-hour mark (DMSO-DMSO, DMSO-TSA, 5’-Ph-IAA-DMSO, and 5’-Ph-IAA-TSA), followed by formaldehyde fixation.

### Cell cycle analysis

For cell cycle analysis, cells were trypsinized and washed using 1xDPBS once and fixed in 90% ethanol at -20°C for at least 24 hours. Fixed cells were washed in 1xDPBS and then resuspended in DPBS containing 2mM MgCl_2_, 0.5mg/ml RNase A (Roche, 10109169001), 1xPI (20xPI in DMSO). The samples were incubated at 20°C for 30min and then analyzed using an LSRII flow cytometry instrument with green channels to monitor DNA contents. FACS data were processed and analyzed using FlowJo v.3. Viability gates using forward and side scatter were set on each sample. DNA content was plotted as a histogram of the PI channel.

### Hi-C experiments

Hi-C for fixed cells was performed as described in our previous study(Liu and Dekker 2022). Briefly, the fixed cells were first lysed to obtain nuclei. After being washed twice with cold NEBuffer 3.1, the nuclei from fixed cells were resuspended in 342ul NEBuffer 3.1 with 0.1% SDS and the tube was gently mixed. The tube was then incubated at 65°C for 10min and put on ice immediately, followed by the addition of 43ul of 10% Triton X-100 and gentle mixing. DpnII (400U) digestion was performed at 37°C overnight with gentle rocking. Once enzyme digestion was completed, the reaction was incubated at 65°C for 15mins to inactivate DpnII. DNA overhanging ends were then filled in with biotin-14-dATP at 23 °C for 4 hours and then ligated with T4 DNA ligase at 16 °C for 4 hours. DNA was treated with proteinase K at 65 °C overnight to remove proteins. Ligation products were purified, fragmented by sonication to an average size of ∼200 bp and size-selected to fragments of 100–350 bp. We then selectively purified biotin-tagged DNA using streptavidin beads before performing end repair, dA-tailing and adaptor addition, using NEBNext Ultra II DNA Library Prep Kit for Illumina (E7645L). Dual indexes were then added by PCR using NEBNext Multiplex Oligos for Illumina (E7600S). The PCR primers were removed from final libraries using AMPure beads. Hi-C libraries were then sequenced using PE150 bases on an Illumina HiSeq 4000 or an Illumina NovaSeq instrument.

### Hi-C for G1 sorted cells

As previously reported(Liu and Dekker 2022) to sort G1 cells for Hi-C analysis, the cells were fixed following the Hi-C protocol using 1% formaldehyde. The fixed cells were washed in 1×PBS and resuspended in PBS containing 2 mM MgCl_2_, 0.1% saponin, 0.5 mg ml−1 RNase A and PI (200 μM stock in dimethylsulfoxide). The samples were incubated at 20 °C for 30 min and then analysed using an BD FACS Aria II flow cytometry instrument using the yellow channels to monitor the DNA content. To avoid obtaining any cells in the S phase, only the cells in the left part of G1 peak were collected. The FACS data were processed and analysed using FlowJo v.3. Viability gates using forward and side scatter were set on each sample. The sorted G1 cells were then used to generate Hi-C libraries as described as above.

### Hi-C and data analysis

Sequencing data of PE100 or PE150 was first trimmed to PE50 using the in-house scripts. All Hi-C PE50 fastq raw sequencing files were mapped onto hg38 human reference genome using distiller-nf mapping pipeline (https://github.com/mirnylab/distiller-nf). After mapping, aligned reads were further processed to remove duplicates (https://github.com/mirnylab/pairtools) to obtain a set of filtered reads defined as valid pairs. Valid pairs were then binned into contact matrices at 10 kb and 100 kb resolutions using cooler50. Intrinsic Hi-C biases were removed using the Iterative Correction and Eigenvector decomposition (ICE) procedure(Imakaev et al. 2012). This was applied to all of the matrices, ignoring the first two diagonals to avoid short-range ligation artifacts at a given resolution, and bins with low coverage were removed using the MADmax filter with default parameters. Contact matrices were stored in ‘.cool’ files and used in downstream analyses.

To identify differences in each condition, Eigenvector 1 values (EV1) were obtained from Hi-C compartment analysis and principal component analysis (PCA) was performed on each condition, CTL vs. KD and DMSO vs. TSA.

For compartment analysis, compartment boundaries were identified in cis using eigen vector decomposition on 50 kb binned data with the cooltools call-compartments function. A and B compartment identities were assigned by gene density tracks such that the more gene-dense regions were labelled A compartments, and the EV1 sign was positive. Changes in compartment type therefore occur at locations where the value of PC1 changes sign. Compartment boundaries were defined at these locations, except for when the sign change occurred within 400 kb of another sign change.

To measure compartmentalization strength, we calculated observed/expected Hi-C matrices for 100 kb binned data, correcting for average distance decay as observed in the *P*(*s*) plots (cooltools compute-expected). We then arranged observed/expected matrix bins according to their EV1 values of the sample without any treatments in each replicate. We aggregated the ordered matrices for each chromosome within a dataset and then divided the aggregate matrix into 50 bins. Strength of A-A and B-B interactions were separately calculated using AA/AB and BB/BA, respectively. The values used for this ratio were determined by calculating the mean value of the 10 bins in each corner of the saddle plot. To compare AA or BB compartment strength changes between CTL and KD, we first calculated the difference of AA or BB interactions between TSA and DMSO in CTL or KD, then normalized to DMSO treatment in each condition. The relative difference is obtained to have the difference of normalized TSA-induced changes between CTL and KD.

For aggregation of loop interactions, the previously identified sets of HCT-116 looping interactions were used(Sanborn et al. 2015; Rao et al. 2017). In total, 3169 looping interactions are on the structurally intact chromosomes of HCT-116. To visualize the looping interactions, we aggregated 20 kb binned data at all loops using Cooltools. We also aggregated 20kb binned data at different sizes of loops, 100kb-500kb, 500kb-1Mb and greater than 1Mb. The size of a loop refers to the distance between the two loop anchors.

For *P*(s) plots and derivatives, the cis reads from the cooler files were used to calculate the contact frequency *(P)* as a function of genomic separation (s) (cooltools). All of the *P*(*s*) curves were normalized using expected cis interactions (cooltools compute-expected) in each dataset. Corresponding derivative plots were calculated from each *P*(s) plot.

To evaluate the different effect of TSA treatment between CTL and KD conditions, the difference in P(s) between DMSO and TSA treatments was first calculated. This difference in CTL conditions were then plotted against KD conditions to determine their deviation from y = x.

To calculate the ratio of TSA-induced P(s) changes in the absence versus the presence of cohesin, the difference in P(s) between DMSO- and TSA-treated samples under each condition was determined. Then the ratio of these differences between cohesin-depleted and cohesin-intact samples was calculated and plotted along genenomic distance.

For interaction aggregation at TAD boundaries, we first calculated observed/expected Hi-C matrices of each sample for 20 kb binned data, correcting for average distance decay as observed in the *P*(s) plots (cooltools compute-expected). We then aggregated the observed/expected Hi-C matrices of each sample at the TAD boundaries that were identified from the sample without any treatments, covering 600kb up and downstream of each boundary, and then generated a pileup heatmap of TAD boundaries for each sample.

To assess the difference in chromatin interactions between DMSO and TSA, log2 of the ratio of TSA/DMSO was performed for each condition, and a heatmap was generated showing increases (red) and decreases (blue) in chromatin interaction frequency for each condition.

For average interactions within TADs, we first identified TAD boundaries by calculating the genome-wide contact insulation score using a 20 kb resolution and boundary calling using the Li threshold (https://github.com/open2c/cooltools). Each TAD region was determined by taking the end position from a TAD boundary and the start position for the subsequent boundary. For each TAD region, the diagonal signal is removed and the number additional bins removed from the diagonal and edges of the TAD are set by the TAD sizes. The intra-TAD intensity is calculated by taking the average contact score of the remaining bins. Outliers in the intra-TAD intensity were removed using the interquartile range. The intra-TAD intensity violin plot was visualized using matplotlib and seaborn.

### Western blot for cohesin components

For each condition, 4 million cells were lysated with 200ul RIPA buffer (Thermo Fisher, #89900) containing protease inhibitor and TurboNuclease (Accelagen, #N0103M). All the protein samples were collected and incubated at 4°C for 10minutes. After spun at 8000g for 3 minutes, the supernatant was transferred to a new tube and 5x sample buffer was added. After mixing, the lysis was boiled at 95°C for 3mins for western blot analysis.

The volume for approximately the same number of cells for each sample was loaded into each lane of a protein gel for separation. Two types of protein gel and buffer were used. To separate small proteins (MW<50KD), NuPAGE 4–12% Bis-Tris protein gels (Thermo Fisher, #NP0322BOX) was used with NuPAGE MOPS SDS Running Buffer (Thermo Fisher, #NP0001). For large proteins (MW > 100kD), NuPAGE 3-8% Tris-Acetate protein gels (Thermo Fisher, #EA03752BOX) were used in NuPAGE Tris-Acetate SDS running buffer (Thermo Fisher, #LA0041). Proteins were transferred to nitrocellulose membranes (Bio-Rad, #1620112) at 30 V for 2 h in 1× transfer buffer (Thermo Fisher, #35040) in the cold room. The membranes were blocked with 5% milk in TBST (20mM Tris-HCl, pH 7.4, 150mM NaCl and 0.1% Tween-20) for 30 minutes at room temperature. The membranes were then incubated with the specified primary antibodies diluted 1:1,000 in TBST overnight at 4°C. The membranes were washed three times with TBST for 10 min at room temperature each, then incubated with secondary antibodies (anti-rabbit IgG HRP-linked, Cell Signaling, 7074) diluted 1:5,000 in TBST for 2 hours at room temperature. The membranes were then washed three times with PBS-T for 10 min each. Then, the membranes were developed and imaged using SuperSignal West Dura Extended Duration Substrate (Thermo, #34076) and Bio-Rad ChemiDoc with Image Lab 6.0.1. ChatGPT (OpenAI, GPT-5.3) was used to assist with language editing; all content was reviewed and revised by the authors.

### Statistics and Reproducibility

No statistical method was used to predetermine sample size. Unless specified in the legends, Western blot analyses have been performed at least three times. All Hi-C and ChIPseq analyses have been performed for two biological replicates.

## Data Availability

Deep-sequencing data that support the findings of this study have been deposited in the Gene Expression Omnibus (GEO) under accession codes GSE******. All other data supporting the findings of this study are available from the corresponding author on reasonable request.

## Code Availability

All programs used in this study are from Open Chromosome Collective (Open2C) and are public available in GitHub: https://github.com/open2c. No other customized codes were developed for this study.

